# Pore-Based RNA Evaluation for Control of Integrity, Sequence, and Errors – Quality Control (PRECISE-QC)

**DOI:** 10.1101/2025.09.20.677417

**Authors:** Yvonne Y. Yee, Dinara Boyko, Bhoomika Pandit, Maksim Royzen, Sara H. Rouhanifard, Meni Wanunu

## Abstract

RNA is at the forefront of therapeutics and gene editing technologies. Yet, RNA synthesis remains expensive and low-yield. Consequently, most oligo manufacturers abstain from synthesizing RNA oligos longer than 60-mers. Solid-phase synthesis is the current standard production method but is often fraught with low coupling yields for canonical nucleotides and even poorer coupling for modifications. This results in high levels of byproducts such as truncations and RNA infidelity. Existing analytical methods can only provide quality control metrics such as RNA length distribution or limited composition information for short oligos. Here, we developed a standard quality control metric using Oxford Nanopore direct RNA sequencing to obtain direct insight into RNA length distribution, sequence, and presence of RNA modification sites. Our pipeline identifies error-prone regions and truncation sites that occur during synthesis. Furthermore, problematic steps in the synthesis are identified and repaired. We show that our platform can produce and assess CRISPR guide RNAs with high-fidelity and higher cleavage activity, and further, that modifications can be reliably detected. We envision that our tool will serve as an integral method for quality control pipelines that assess the integrity and accuracy of synthetic RNAs and guide the improved synthesis and yield of synthesized RNAs.

**Graphical Abstract:** 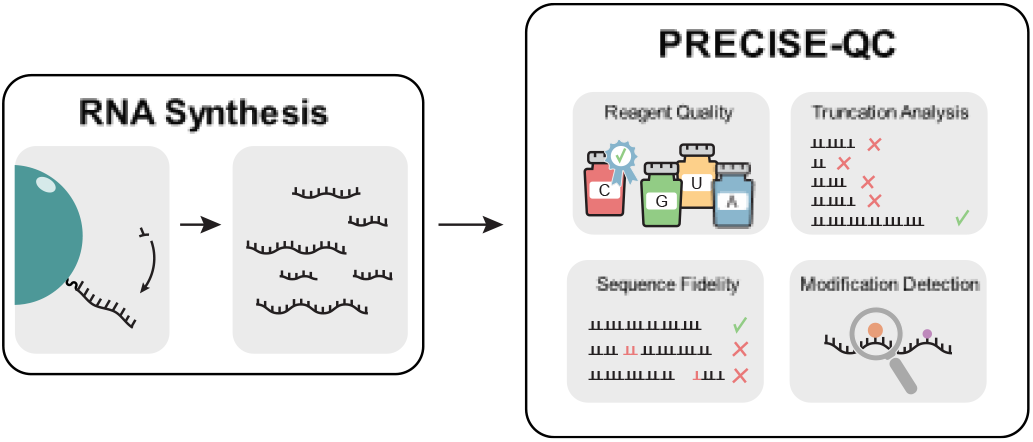

## Introduction

The demand for RNA manufacturing is rapidly increasing for various applications ranging from pharmaceuticals to agriculture. RNA oligonucleotides can selectively bind to target RNA and DNA to modulate gene expression and can encode antigenic proteins to elicit immune responses [1]. These properties have enabled breakthroughs in disease treatment, vaccinations, as well as crop improvement [2–5]. Incorporating chemically modified bases further enhances RNA performance by increasing binding affinity, improving stability, and providing improved nuclease resistance [6–8]. The high demand for large-scale RNA manufacturing has led to a current synthetic RNA market valued at approximately $1 billion [9]. Despite this growth, RNA synthesis remains inefficient and costly, posing significant challenges for large-scale production [10–12].

The current gold standard for RNA oligonucleotide synthesis is solid-phase synthesis using phosphoramidite chemistry. This method involves anchoring the initial 3’ nucleotide to controlled pore glass (CPG) beads, followed by a four-step cycle comprised of detritylation, coupling, capping, and oxidation steps to synthesize RNA from 3’ to 5’ one nucleotide at a time (Figure 1A). In between steps are capping reactions and washes to prevent propagation of failure sequences, i.e., sequences that do not match the reference [13]. Some industrial processes, however, are known to eliminate the capping step [11]. Natural phosphoramidites can be replaced with modified phosphoramidites to specifically incorporate base modifications within the sequence [14,15].

**Figure 1.**
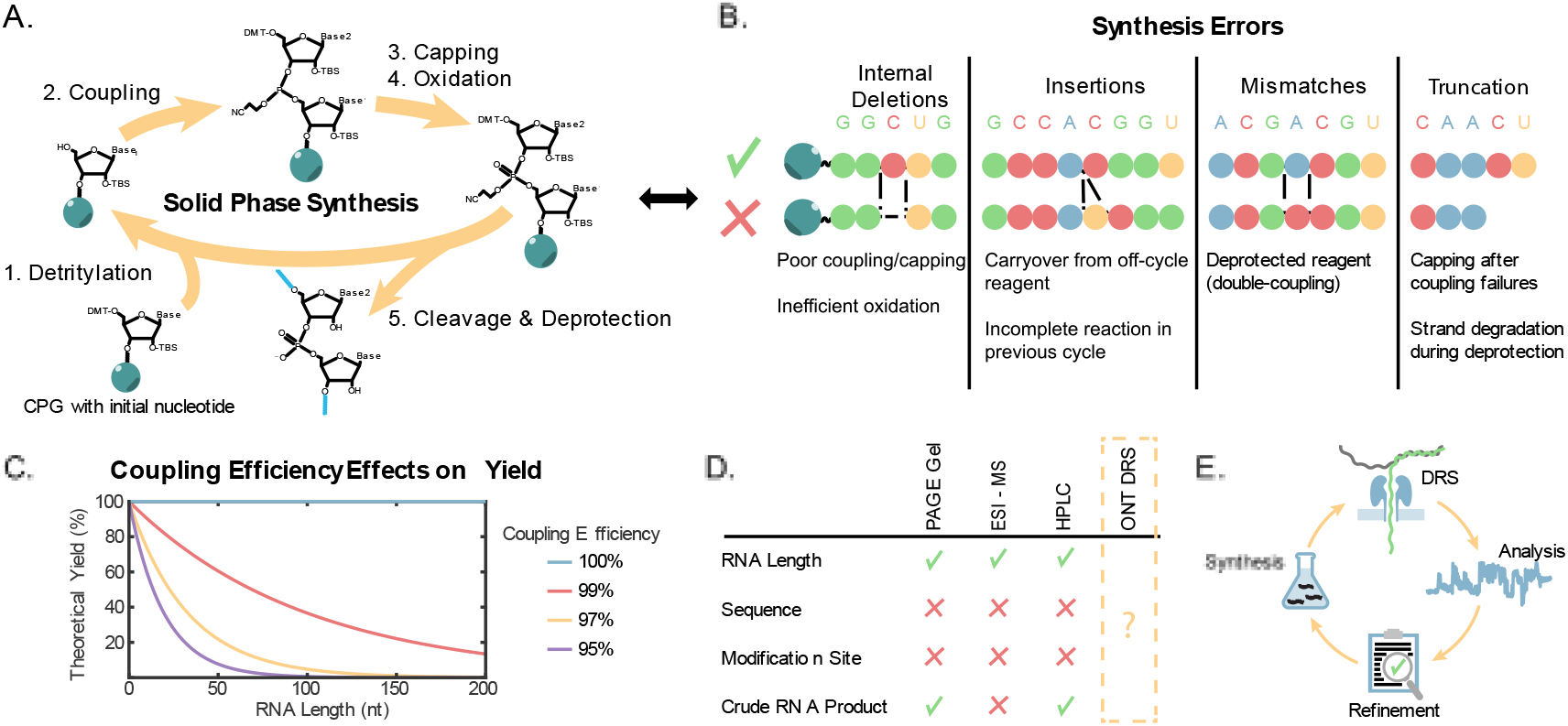
Overview of RNA solid-phase synthesis and quality control methods **(A)** Schematic of RNA solid-phase synthesis steps for extending RNA one base at a time starting from the initial 3’ nucleotide on the CPG bead. **(B)** List of error types and causes that can occur during solid-phase synthesis. **(C)** Plot of theoretical yield vs. RNA length based on the effects of coupling efficiency. **(D)** A comparison chart of quality control tools, showcasing the benefits of ONT DRS. **(E)** The suggested pipeline on how to integrate ONT DRS as a standard quality control for RNA synthesis.

The yield of the target RNA product is the quantity of RNA synthesized with the expected length but assumes that all full-length RNA contains the correct sequence. Realistically, the RNA product can contain impurities such as failure sequences that managed to slip through the screening undetected due to having the correct length. These failure sequences are generated and accumulated at every cycle due to incomplete coupling. Other mishaps that are less frequently spoken about but can nonetheless occur are due to partial reagent degradation, inefficient washing steps, double-coupling, or subunit misincorporation. These synthetic flaws lead to deletions, insertions, and/or mismatches occurring within the sequence (Figure 1B). Ideally, synthesis operating at 100% coupling efficiency will not contain any failure sequences. However, even a 1% drop can be detrimental to RNA yield and purity (Figure 1C). The accumulating errors that occur each cycle is why long RNA (>55 nt) is notoriously difficult to synthesize and purify. Factors such as instrumentation, reagent quality and RNA’s complexity can all play a role in diminishing synthesis operations. Synthesis problems are further exacerbated when incorporating RNA modifications due to increased structural complexity leading to significantly lower coupling yields [16]. A preventative measure to reduce propagation of failure sequences is capping reactions that occur in between steps, leading to truncated byproducts. Evaluating full-length failure sequences can potentially help pinpoint synthesis problems such as reagent quality and reaction parameters in order to improve coupling efficiency.

Due to the high cost of RNA production, optimizing synthesis and quality control (QC) methods is essential. Current QC methods including gel electrophoresis, electrospray ionization mass spectrometry (ESI-MS), and high-performance liquid chromatography (HPLC) have significant limitations (Figure 1C). Gel electrophoresis and HPLC can assess length distribution but provide no sequence information [17]. ESI-MS often used with HPLC, can measure nucleotide mass composition, confirming if the correct base distribution is incorporated into the sequence. Yet, it cannot fully confirm if the product contains the correct sequence. ESI-MS analysis is further complicated by the difficulty of desalting long RNAs. To confirm the sequence fidelity, methods such as Illumina-based RNA sequencing are done which are short read and provide no information about modifications and length distribution of product. This results in ∼90% purity of many RNA-based therapeutics currently in the market with majority of the impurities being stereoisomers [12,18]. There is an unmet gap in the field for a method that would provide 1. length distributions, 2. sequence analysis, 3. base composition, and 4. total yield of RNA synthesis.

Direct RNA sequencing (DRS) by Oxford Nanopore Technologies (ONT) offers a promising alternative that complements and surpasses conventional QC tools (Figure 1D). ONT DRS is the only platform capable of sequencing RNA directly without the need for reverse transcription and PCR which erases RNA modification and amplifies the sequence [19]. During sequencing, an applied voltage drives the RNA strand through the nanopore, generating electrical current traces that reflect the identify of each nucleotide. Therefore, current traces can be processed to provide the RNA sequence, including sequences containing certain base modifications.

In this study, we show how ONT DRS can be incorporated into the RNA synthesis pipeline as a new quality control standard that will lead to accelerating low-cost, high-quality RNA production (Figure 1E). This method, Pore-Based RNA Evaluation for Control of Integrity, Sequence, and Errors – Quality Control (PRECISE-QC), enables direct sequencing of crude RNA to analyze in silico purified full length sequences, circumventing the need for purification steps during quality analysis. PRECISE-QC provides more reliable length distribution and accurate yield value since it is capable of distinguishing between failed and successful sequences that contain the expected length. Additionally, its capacity to detect select base modifications allows for rigorous inspection of chemically modified RNA. These insights can be used to iteratively refine synthesis protocols and improve product quality.

## Materials and Methods

### Solid-Phase Synthesis

All oligonucleotide solid phase syntheses were done on a 1.0 μmol scale using the Oligo-800 synthesizer (Azco Biotech, Oceanside, CA, USA). Solid phase syntheses were performed on control-pore glass (CPG-1000) purchased from Glen Research (Sterling, VA, USA). Other oligonucleotide solid phase synthesis reagents were obtained from ChemGenes Corporation (Wilmington, MA, USA). Phosphoramidites (TBDMS as the 2′-OH protecting group): rA was N-Bz protected, rC was N-Ac protected and rG was N-iBu protected. A, C, G, U phosphoramidites were dissolved in anhydrous acetonitrile (0.07 M) directly before use. m^6^A and Y phosphoramidites were dissolved in anhydrous acetonitrile (0.15 M) directly before use. Coupling step was done using 5-ethylthio-1H-tetrazole solution (0.25 M) in acetonitrile for 12 min. 5′-detritylation step was done using 3% trichloroacetic acid in CH_2_Cl_2_. Oxidation step was done using I_2_ (0.02 M) in THF/pyridine/H_2_O solution.

### CRISPR-Cas9 in vitro Cleavage Assay

eGFP-N1 plasmid DNA (10 U/μL, 1 μL, NEB, R3510L) was diluted with water (16.87μL) and NEB buffer 3.1 (10x, 2 μL). The plasmid was linearized directly prior to CRISPR with DraIII-HF (10 U/μL, 1 μL, NEB, R3510L). For the Cas9-mediated DNA cleavage assay, sgRNA (300 nM, 5 μL), Cas9 (1 μM, 0.3 μL, NEB, M0386S), Cas9 buffer (10x, 1 μL, NEB), linearized plasmid (20 nM, 1.5 μL) and MQ H2O (2.2 μL) were mixed (final volume = 10 μL) and incubated for 1h, 2h and 4h at 37 °C. CRISPR experiments were terminated by the addition of proteinase K (20 mg/mL, 0.5 μL) for 1 h at 37 °C. The reaction (10 μL) was mixed with blue loading buffer (6x, 2 μL, NEB, B7703S) and loaded on a 1% agarose stained with ethidium bromide (1x TBE running buffer).

### Synthesis of RNA Adapter

To produce 5’ adapters for ligation to synthetic RNAs we created an RNA template that is based from a human genome. Specifically, a gene block containing the T7 promoter site upstream from a partial sequence of either the human MRPS14 or PSMB2 gene as well as forward and reverse primers (Table S1) were purchased from Integrated DNA Technologies (IDT). The gene block and primers were used to amplify 117 nt DNA templates using *Taq* PCR kit (New England Biolabs (NEB) E5000S). During PCR, 0.5 ng of gene block was used with 0.2 uM primers. The template was then purified using the Monarch Spin PCR & DNA Cleanup kit (5 μg) (NEB T1130S) and eluted in 15 μL nuclease-free (NF) water.

The template was then used to synthesize RNA through *in vitro* transcription (IVT) using the HiScribe T7 Quick High Yield RNA Synthesis kit (NEB E2050S). During IVT, 1 μg of template was used and the sample was incubated at 37°C for 2 h. After IVT, the sample was incubated at 37°C for 15 min with 4 U DNase I (RNase-free) to digest all DNA. Then, the sample was purified using the Monarch Spin RNA Cleanup Kit (500 μg) (NEB T2050S) and eluted in 20 μL NF water.

### Ligation of RNA Adapter and sgRNA

The sgRNA samples were phosphorylated on the 5’ end using T4 Polynucleotide Kinase (NEB M0201S). 10 U of kinase were used to phosphorylate 300 pmol of RNA at 37°C for 1 h followed by heat inactivation at 65°C for 20 min. The phosphorylated sgRNA was then mixed with adapter RNA and ligated together using T4 RNA Ligase 1 (NEB M0204S). For each ligation sample, there was 15% PEG 8000, 10% DMSO, 1 mM ATP, 20 U NEB murine RNase inhibitor, and 20 U T4 RNA ligase 1. The unmodified GFP sgRNA was mixed with the MRPS14 adapter RNA at a 1:4 molar ratio. All other ligated samples were mixed at 2:1 sgRNA to adapter molar ratios. The EP300 Nm modified sgRNA was the only sample that used the PSMB2 adapter. Samples were incubated at 25°C for 2 h. They were then purified using the Monarch Spin RNA Cleanup Kit (500 μg) and eluted in 20 μL NF water.

### RNA Sequencing

Poly(A) tails were added to the ligated RNA adapter-sgRNA samples using the *E. coli* Poly(A) Polymerase kit (NEB M0276S). 10 U polymerase was used to add a poly(A) tail to the entire ligated sample. The sample was incubated at 37°C for 10 min. RNAClean XP beads (Beckman Coulter A63987) were used to purify the poly(A) tailed samples and were eluted in 12 μL NF water. 300 ng of the poly(A) tailed samples (first synthesis unmodified GFP sgRNA, optimized synthesis unmodified GFP sgRNA, and optimized synthesis modified GFP sgRNA) were used to prepare libraries following the Nanopore Direct RNA sequencing kit documented protocol (SQK-RNA004) (DRS_9195_v4_revF_11Dec2024) and sequenced using minion (FLO-MIN004RA) flow cells. Alternatively, the 150 ng each of the poly(A) tailed Nm modified GFP and EP300 sgRNA were combined for a total of 300 ng. The same Nanopore Direct RNA sequencing kit protocol was used for the Nm modified sgRNAs followed by sequencing using a PromethION (FLO-PRO004RA) flow cell.

### Basecalling, Filtering, and Alignment for in silico Purified Full-Length Reads

Our PRECISE-QC pipeline is described in the GitHub page (see Software Availability section for link). Briefly, Pod5 files containing the ionic current of sequenced reads were basecalled using Dorado V1.0.0 and applied the rna004_130bps_sup@v5.2.0 model [20]. During basecalling, each dataset was set to detect pseudouridine, m6A, and 2’O methylations (parameters ‘--modified-bases pseU_2OmeU m5C_2OmeC 2OmeG inosine_m6A_2OmeA --emit-moves’). Basecalled reads were then aligned to the sgRNA reference sequence using BWA-MEM V0.7.15 (parameters ‘-t 1 -w 13 -k 6 -x ont2d’) [21]. Reads were then filtered for primary alignments and soft-clippings were removed to retrieve the length distribution using samtools V1/21. Full length reads (95 nt – 105 nt) were filtered and analyzed for their indel levels by base, motif, and full sequence basis using pysamstats V0.15.2 (parameters --type variation) to generate a per-nucleotide error profile [22,23].

### Filtering and Alignment for Truncated Reads

The basecalled reads were then filtered using Porechop V0.2.4 to only collect 5’ adapter containing reads (parameters ‘--barcode_diff 1 --barcode_threshold 74) [24]. These reads were then aligned and filtered the same as above to retrieve the length distribution. The truncated reads were then filtered by size for easier analysis using Integrative Genomics Viewer (IGV).

### Modification Signal Assessment

Nanopore current signal was aligned to the reference sequence with Uncalled4 (alignment parameters ‘--min-aln-length 50’, alignment in event align format parameters ‘--eventalign-out --eventalign-flags print-read-names,signal-index,samples’). Uncalled4 utilizes basecall-guided dynamic time warping to map ionic current to basecalled reads in eventalign format [25]. The output of mapping is a table saved in tsv format with current signal values for each kmer. The signal values were then visualized using numpy library numpy.linearspace.

### Overall and Base Synthesis Accuracy Definitions

Using pysamstats variation parameter, we can pull the number of matches, mismatches, deletions, and insertions occurring at all positions along the sequence reference for all full-length reads. Using this dataset, we plotted the frequency of total error rates along the reference. Overall synthesis accuracy is the percentage of sites containing total error rates below the 5% error threshold. Base synthesis accuracy is the percentage of specific base sites containing total error rates below the 5% error threshold.

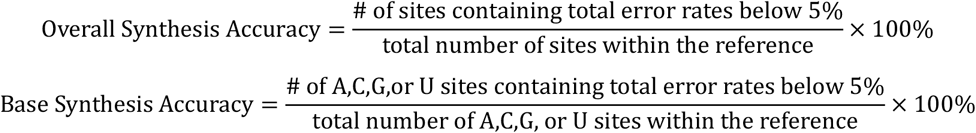

### Average Read Accuracy and Error Rate Definitions

The number of matches, mismatches, insertions, and deletions for each read is determined by parsing through the CIGAR string and MD tag within the .sam files containing full-length reads. The read accuracy is the summed matches divided by the summed matches, mismatches, insertions, and deletions within a read [23,26]. The average read accuracy is the mean of all full-length read accuracies. Similarly, the mismatch, insertion, and deletion rates are the summed error types over the summed matches and errors within a read. The average error rate is the mean of all full-length read error rates.

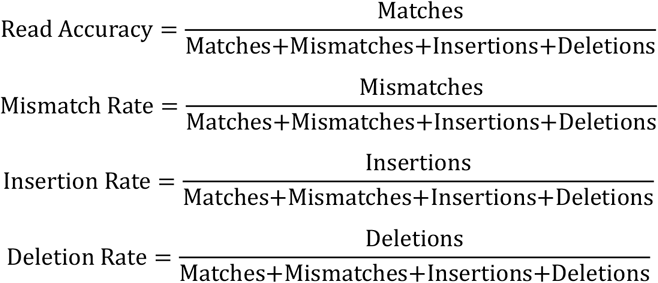

## Results

To validate our PRECISE-QC approach, we synthesized a 101-nt single guide RNA (sgRNA) using solid-phase synthesis. Accurate sgRNA sequence is crucial for gene editing using CRISPR-Cas9; sgRNA guide region must be synthesized correctly for selective DNA targeting while stem loop fidelity is relevant for stabilizing the sgRNA-Cas9 complex (Figure 2A) [27]. Deletions, insertions, and/or mismatches occurring within the sequence could potentially result in off-target binding. We first sequenced an unmodified (i.e., using only canonical nucleotides) sgRNA targeting GFP using ONT DRS.

**Figure 2.**
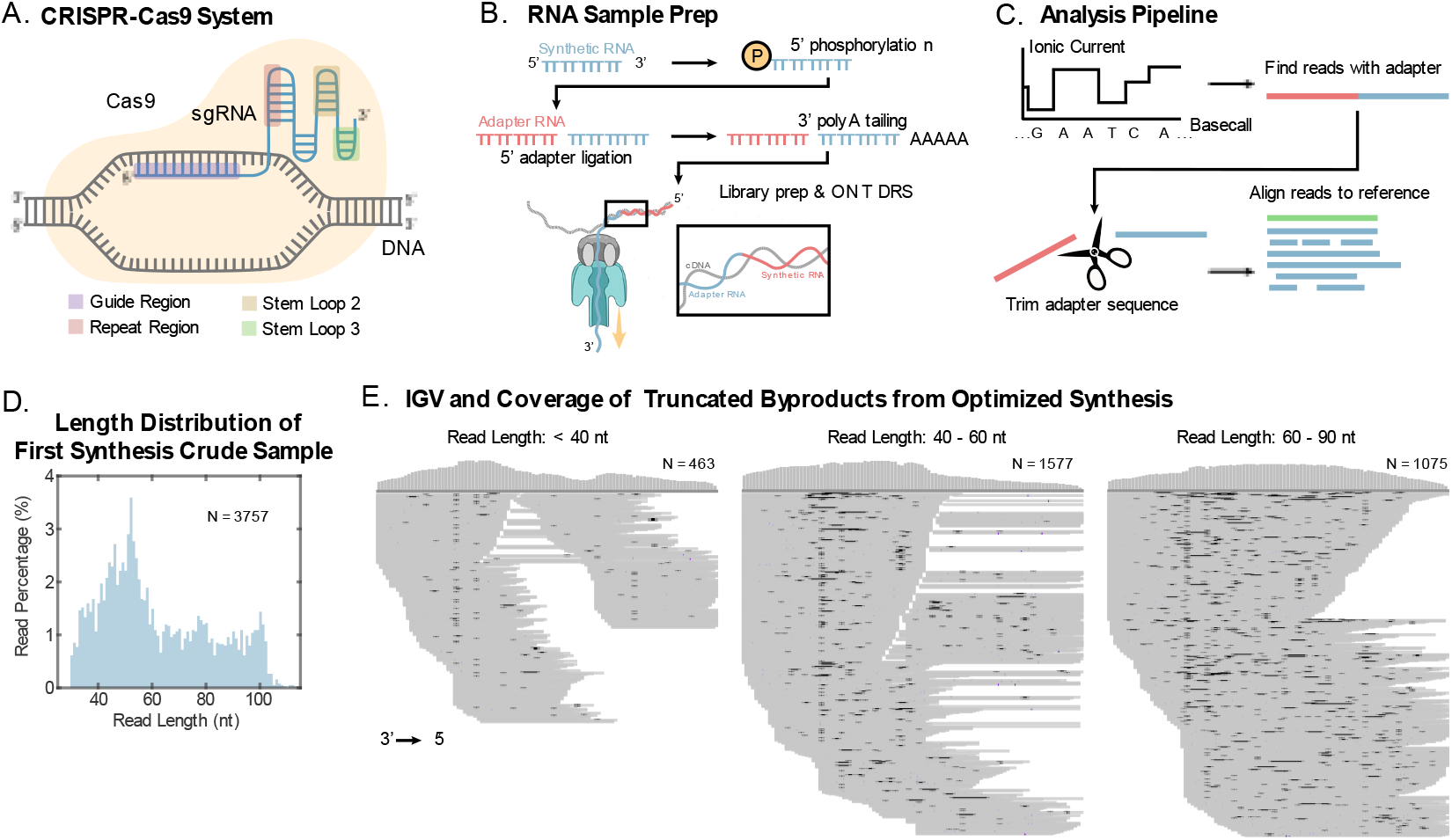
RNA synthesis sample preparation and analysis of truncated byproducts. **(A)** CRISPR-Cas9 system with sequence relevant regions of a single-guide RNA (sgRNA) highlighted. **(B)** RNA sample preparation schematic of ligating a 5’ adapter to the synthetic RNA to allow for full coverage sequencing of synthetic RNA crude sample using ONT DRS. **(C)** Computational pipeline to isolate adapter containing reads for further analysis. **(D)** Length distributions of first synthesis unmodified GFP sgRNA crude sample. **(E)** IGV and coverage plots of length subsets (<40 nt, 40 – 60 nt, and 60 – 90 nt) from first synthesis crude sample. Each gray bar on an IGV plot correlates to a single read. Rows that show a break in the gray bar equates to multiple reads placed on the same row. Peaks in the coverage show that many reads align to that particular region in the reference.

During sequencing, ONT DRS utilizes enzyme motor ratcheting to drive RNA through the nanopore in a base-by-base manner. This discretizes the signals from RNA k-mers within the pore constriction, thereby increasing the RNA resolution and improving read quality. Because the motor is positioned above the nanopore, the 5’ end of the RNA is often dislodged before the last 10-15 nucleotides can be read. To recover this region, we applied a commonly used technique to ligate a 5′ adapter to the synthetic RNA, enabling complete sequence capture in every adapter containing synthetic RNA (Figure 2B) [28,29].

### Truncation Analysis

Both PAGE gels and HPLC can be used to show the lengths of byproducts generated during solid-phase synthesis. However, they lack the ability to determine where truncation, fragmentation, and RNA degradation are likely to occur. ONT DRS can provide a more accurate length distribution compared to the other QC methods by relying on read count instead of absorbance and can also provide insight on where truncations or degradation are likely to occur during synthesis.

As RNA molecules are enzymatically ratcheted through the pore, they generate current traces which are then parsed and converted to a corresponding RNA sequence using basecalling software. To ensure truncation patterns reflect synthesis rather than sequencing artifacts, we first filtered reads to include only those containing the 5′ adapter (Figure 2C). This allowed us to accurately determine read lengths and analyze synthetic byproducts. We defined full-length reads as those between 95–105 nt and plotted the length distribution of the first synthesis crude sample (Figure 2D).

To investigate where truncation and degradation occurred during synthesis, we analyzed read alignments grouped by length: less than 40 nt, 40–60 nt, and 60–90 nt. Integrative Genomics Viewer (IGV), a visualization tool of sequences aligned to a reference, revealed distinct alignment patterns for each group (Figure 2E). The <40 nt subset shows two peaks in the coverage: one on the 3’ half and the other on the 5’ half of the reference sequence. Within the 40 – 60 nt subset, most reads were aligned to the 3’ end. Lastly, the 60 – 90 nt subset shows a centered single peak coverage.

### Overall Errors of Full-Length Reads

Alternative QC methods determine product yield by the quantity of full-length RNA. However, they fail to distinguish the sequence fidelity of the product, leading to stereoisomer impurities [12,18]. Our approach to identify full-length RNA sequences and quantitate their accuracy to the reference can ensure product purity during production. To enrich for full-length products, we filtered aligned reads by length, retaining only those with ≥94% coverage. These reads were used to assess the integrity and sequence identity of the synthetic RNA.

The sequencing read accuracy refers to how well a sequence is correctly identified compared to the true sequence and is influenced by several factors such as the sequencing technology, read length, and library preparation quality [30]. It is essential for sequencing systems to generate high read accuracies and low error rates for reliable interpretation of sequencing data. For ONT DRS, the average sequencing read accuracy is around 90% while the mismatch and insertion rates are within 1-2% and the deletion rates often appear higher around 5% [23]. All sgRNA samples that were sequenced showed above 90% average read accuracy, suggesting that further error analysis is likely caused by synthesis (Table 1). The average error rates were also within reasonable parameters; mismatch and insertion rates were below 2% while deletion rates were around 5% or lower.

**Table 1.**
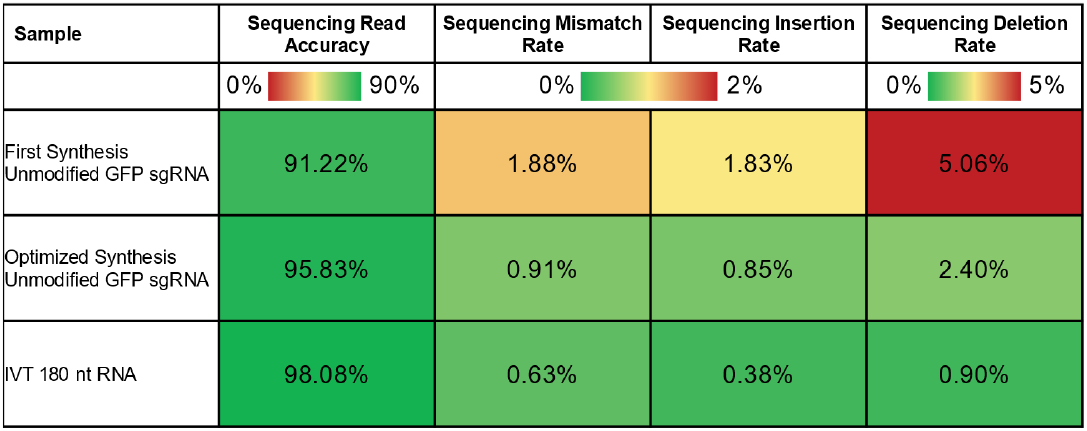
Summary of RNA sequencing quality. The sequencing read accuracy is on average how well full-length reads align to the reference. The passing sequencing read accuracy is above 90%. Similarly, the sequencing mismatch, insertion, and deletion rates are on average the rate of error type occurring per read. Passing sequencing mismatch and insertion rates are below 2%. Passing sequencing deletion scores are below 5%. Lastly, the perfect full-length read is the percentage of full-length reads that do not contain any error types within the sequence. Passing full-length read percentage is above 5%.

| Sample | Sequencing Read Accuracy | Sequencing Mismatch Rate | Sequencing Insertion Rate | Sequencing Deletion Rate |
| --- | --- | --- | --- | --- |
|  | 0% 90% | 0% 2% |  | 0% 5% |
| First Synthesis<br>Unmodified GFP sgRNA | 91.22% | 1.88% | 1.83% | 5.06% |
| Optimized Synthesis<br>Unmodified GFP sgRNA | 95.83% | 0.91% | 0.85% | 2.40% |
| IVT 180 nt RNA | 98.08% | 0.63% | 0.38% | 0.90% |

To define synthesis error rates, we included a 180 nt *in vitro* transcribed (IVT) RNA to set our error threshold (Table S1). Sites along the reference that show error rates below the error threshold are considered correctly incorporated during synthesis. IVT is known to synthesize highly accurate RNA with error rates as low as 10^−5^ to 10^−6^ events per base [31]. To retain above 90% synthesis accuracy for the IVT RNA, we set our error threshold to 5% (Table 2, Figure S1,2A).

**Table 2.**
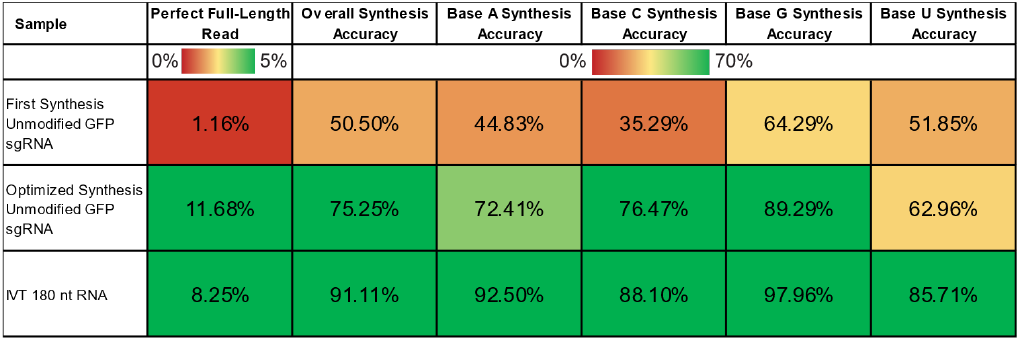
Summary of RNA synthesis quality. Overall and base synthesis accuracy rates provides insight to synthesis efficiency while sequencing read accuracy and error (mismatch, insertion, deletion) rates provides insight on sequencing accuracy. Overall synthesis accuracy is the percentage of sites within full-length reads that show error frequency levels falling within the 5% error threshold. Base synthesis accuracy is the percentage of sites below the 5% error threshold based on the reference base (A, C, G, U). Passing overall and base synthesis accuracy scores (green) are percentages above 70% while failing scores (red) are below.

| Sample | Perfect Full-Length Read | Overall Synthesis Accuracy | Base A Synthesis Accuracy | Base C Synthesis Accuracy | Base G Synthesis Accuracy | Base U Synthesis Accuracy |
| --- | --- | --- | --- | --- | --- | --- |
|  | 0% 5% | 0% 70% |  |  |  |  |
| First Synthesis Unmodified GFP sgRNA | 1.16% | 50.50% | 44.83% | 35.29% | 64.29% | 51.85% |
| Optimized Synthesis Unmodified GFP sgRNA | 11.68% | 75.25% | 72.41% | 76.47% | 89.29% | 62.96% |
| IVT 180 nt RNA | 8.25% | 91.11% | 92.50% | 88.10% | 97.96% | 85.71% |

We analyzed the unmodified GFP sgRNA to identify regions prone to synthesis-related deletions, insertions, and mismatches according to sequence alignment to the reference (Figure 2A top). Only 50.50% (51 out of 101) positions aligned to the reference passed the error threshold in the initial synthesis (Table 2). Two regions stood out with notably high error rates. The repeat/guide region averaged 11.79 ± 5.36% total error, peaking at 23.86%, with mismatches as the dominant error type (6.46 ± 3.18%). Deletion rate was relatively high within the first U of the 3’-UUUU-5’ in the repeat/guide region. The stem loop 2 region showed average error and deletion rates of 9.02 ± 5.97% and 4.13 ± 5.89%, respectively, with deletions concentrated at the first A in the 3′-AAAAA-5′ motif. The nucleotide prior to both long homopolymers (>3 nt) within the problematic regions (G before 3’-AAAAA-5’ and A before 3’-UUUU-5’) showed higher than average insertion rates.

The unmodified GFP sgRNA was resynthesized, accounting for the error prone regions discovered in the initial sequencing run. The resulting error profile showed significantly less errors (Figure 3A middle). 75.25% positions were below the 5% error threshold, a nearly 25% increase compared to the first synthesis. The stem loop 2 average error rate was 6.75 ± 5.03% and average deletion rate was 2.80 ± 4.84%. The error rate dropped by 2.27% while the deletion rate dropped by 1.33% compared to the first synthesis. Similarly, the repeat/guide region average error rate and average mismatch rate dropped by 4.57% and 2.49%, respectively. The average error rate was 7.22 ± 3.75% and the average mismatch rate was 3.97 ± 2.31%.

**Figure 3.**
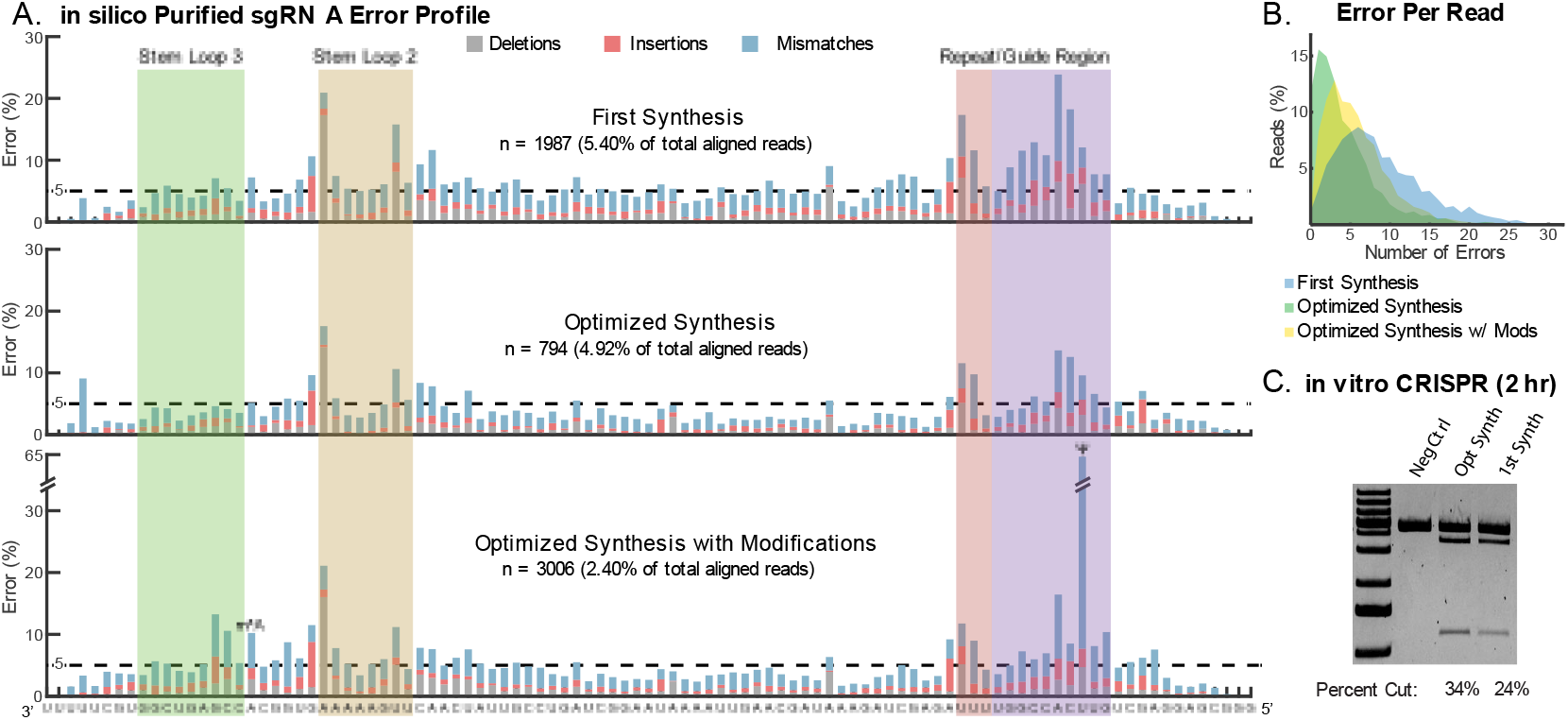
Full-length RNA error analysis. **(A)** Error profile of in silico purified sgRNA before and after quality consultation. **(B)** Plot of number of errors that occur per full-length read. **(C)** in vitro CRISPR cleavage assay comparing percent of cleavage that occurs to eGFP-N1 linearized plasmid after 2 hours with sgRNA from first synthesis vs. optimized synthesis. Negative control is CRISPR reaction without sgRNA.

We also synthesized a GFP sgRNA with pseudouridine (Ψ) and N6-methyladenosine (m^6^A) using the optimized process (Figure 3A, bottom). As expected, the Ψ site—often miscalled as C in DRS—was excluded from error calculations [32–34]. Despite the added synthetic complexity, 61.00% of positions passed the error threshold, outperforming the initial synthesis. Stem loop 2 and repeat/guide error rates were 8.03 ± 5.80% (deletion: 3.57 ± 5.27%) and 8.09 ± 3.57% (mismatch: 4.69 ± 2.49%), respectively— intermediate between the initial and optimized unmodified versions.

Across the full-length reads, mismatches were the dominant error type, averaging 3.24 ± 2.09% in the initial synthesis, and improving to 2.25 ± 1.60% and 2.74 ± 1.72% in the optimized unmodified and modified samples, respectively. Deletions followed, with average rates of 1.57 ± 2.15%, 1.06 ± 1.63%, and 1.26 ± 1.79%, respectively. Insertions were minimal outside of homopolymer contexts. Per-read error summaries showed a clear improvement: the initial synthesis yielded a median of 6 errors per read (max 32), while the unmodified and modified optimized versions dropped to 1 error per read (max 24) and 3 errors per read (max 28), respectively (Figure 3B). A CRISPR-Cas9 in vitro cleavage assay further confirmed functional gains—the optimized sgRNA achieved ∼10% greater target DNA cleavage after 2 hours (Figure 3C), supporting the effectiveness of our synthesis refinement strategy.

### Error Relation to Single Nucleotides

We analyzed synthesis errors that occurred immediately after correctly incorporated base to assess phosphoramidite monomer quality and reagent-driven fidelity during solid-phase RNA synthesis (Figure 4A). During the first synthesis, mismatches were the most common error type, with insertion and deletion rates relatively consistent across bases except for guanine (Figure 4B). The overall error rate remained below 30% for each base and similar patterns were observed for the unmodified and modified sgRNAs— suggesting comparable performance across monomer stocks.

**Figure 4.**
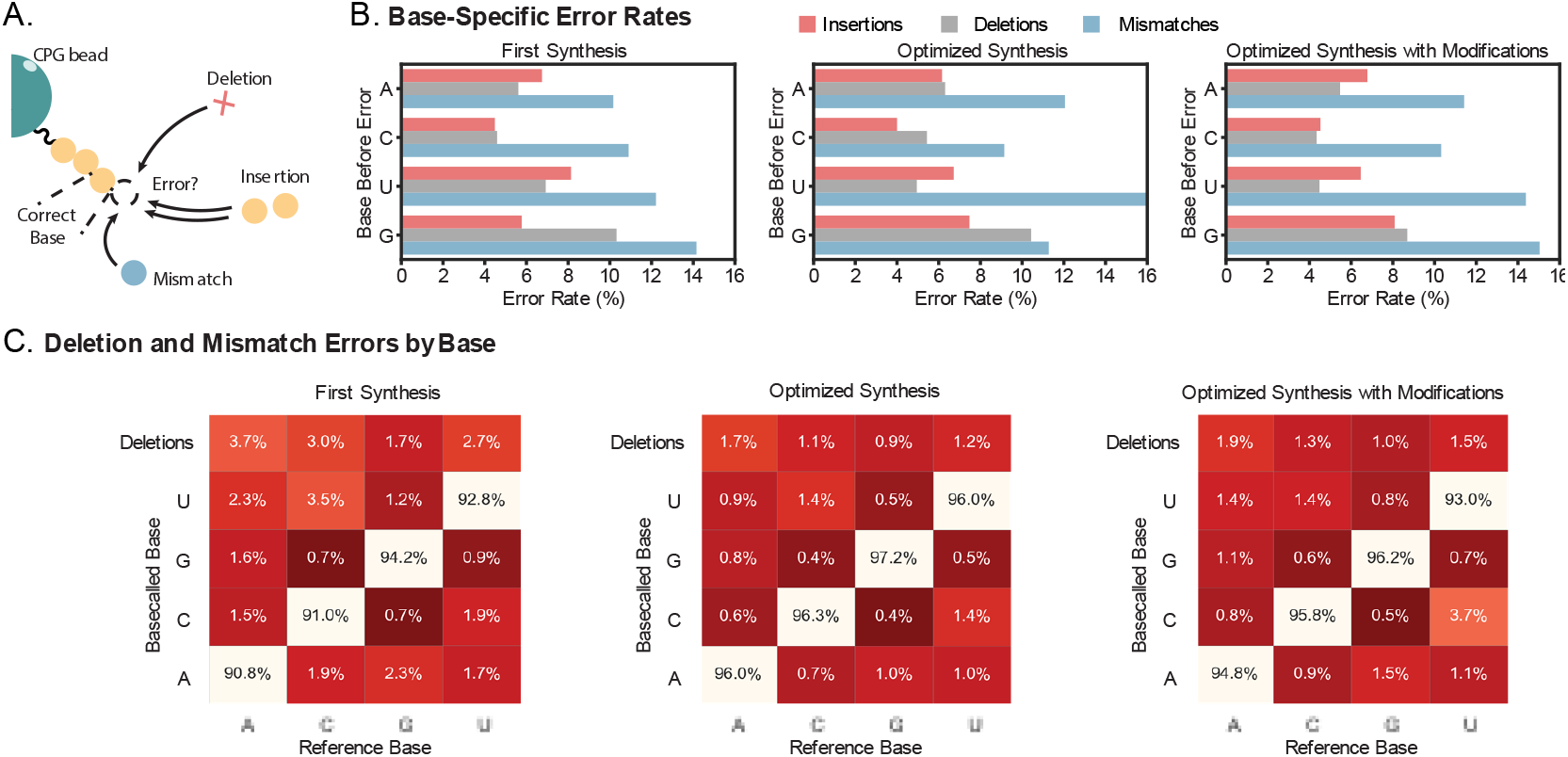
Error rates by base type. **(A)** Schematic of which error is counted during error rate analysis. If a misincorporated base occurs after a correct base, it is counted in the error rates during analysis. **(B)** Bar plots of types of errors that occur by base type. **(C)** Confusion matrices of deletion and mismatch rates that occur by base type.

Confusion matrices were generated to determine what mismatches and deletions occur for each base (Figure 4C). Mismatches between U-C, C-U, and G-A were slightly elevated. Notably, A mismatches were predominantly U during the first synthesis. After synthesis optimization, A-U mismatches were reduced to similar levels as A-G. The optimized synthesis also lead to > 2% mismatches for each type of mismatch while the first synthesis had mismatch rates as high as 3.5%. The modified sgRNA prepared with the optimized synthesis showed similar results as the unmodified except for U-C mismatch that was influenced by the Ψ site.

### Modification Detection Using ONT DRS

Currently, the ONT basecalling software, Dorado V1.0.0, can distinguish a few types of RNA base modifications (5-methylcytosine, m^6^A, inosine, Ψ, and 2’-O-methylation of all four canonical nucleotides) [20]. The 9-mer space is the channel within the nanopore that RNA translocates through (Figure 5A). The current signals generated during translocation are dependent on the 9-mer sequence that currently occupies the 9-mer space [35]. Within our studies, we utilized Dorado’s modification detection on the Ψ and m^6^A modified GFP sgRNA as well as a 2’O methylated (Nm) sgRNA (Table S1).

**Figure 5.**
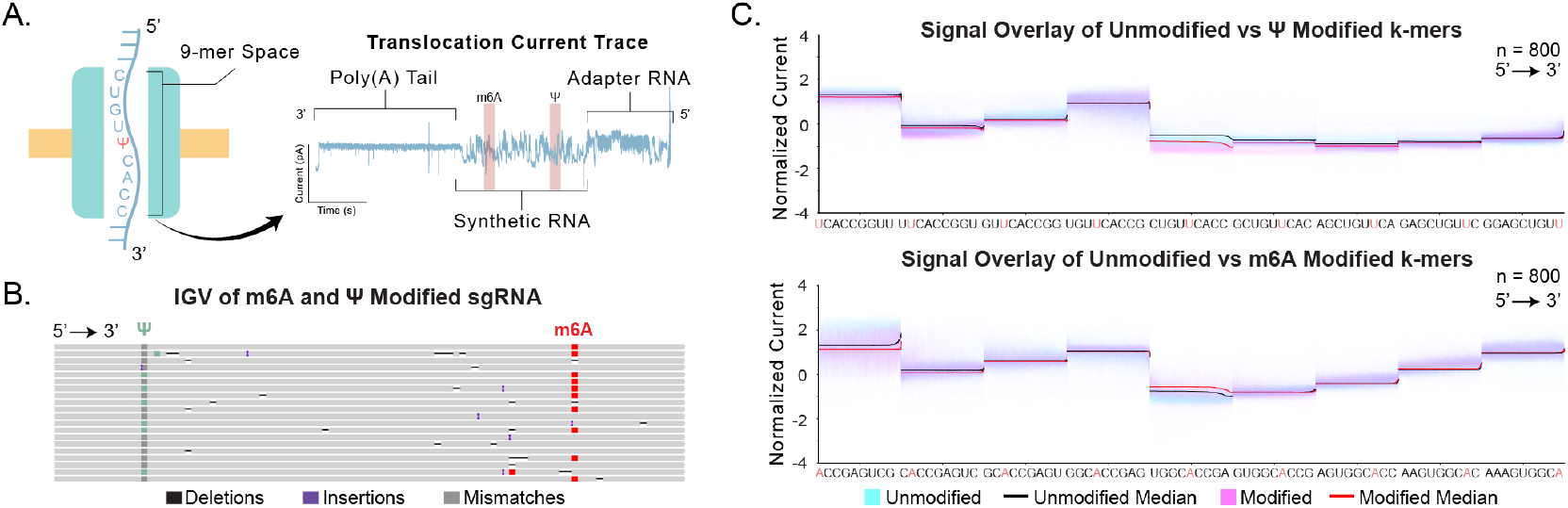
Modification detection using ONT DRS. **(A)** Schematic of current levels caused by the 9-mer sequence that is present within the 9-mer space during translocation through the nanopore. **(B)** IGV plot showing m^6^A and Ψ detection using Dorado modification base calling. **(C)** Current signals of 9-mers containing Ψ at different positions compared to the unmodified sequence (top) as well as 9-mers containing m^6^A (bottom).

The bases flagged as modifications from Dorado basecalling can be visualized using IGV (Figure 5B). In IGV, the highlighted regions of the reads demonstrate the alignment between predicted modification sites and the synthetic reference. The IGV results showed that Ψ and m^6^A are at the expected position, showing that they were properly incorporated during RNA synthesis. If we were to apply a 75% base modification likelihood threshold on IGV, the Ψ site showed 24.93% of full-length reads averaging a 99% likelihood of containing Ψ. For the m^6^A site, there were 65.01% of reads with 92% average likelihood.

The latest release of Dorado V1.0.0 introduced a new model for detecting Nm. However, when we sequenced the Nm sgRNA, the model failed to distinguish between canonical and modified nucleotides (Figure S5). Instead we analyzed the ionic current signal to investigate whether modified bases produce a defined alteration in the current. Previous studies have shown that RNA modifications cause deviations in the raw ionic current signals [25,33]. We examined the ionic current of the sequence containing Ψ and m^6^A (Figure 5C). Both sites demonstrated a clear difference in the current level compared to the unmodified sequence. Additionally, we explored the Nm sgRNA where current levels frequently demonstrated shifts in current at the modified sites (Figure S5). It confirms that the approach to studying synthetic RNA through the raw signal is perceptive enough to control the incorporation of modified bases into synthetic RNA.

## Discussion

Here, we present PRECISE-QC, a quality control pipeline to assess sequence fidelity to refine long RNA synthesis. As the desired RNA length grows longer, more failure sequences appear during the synthesis that are challenging to identify and separate with current quality control and purification methods. In our pipeline, the crude sample produced from solid-phase synthesis is sequenced using ONT DRS to identify potential error prone regions within in silico purified full-length reads, confirm reagent quality, and investigate causes of truncated byproducts.

The error analysis of full-length GFP sgRNA showed improvements after one iteration of our pipeline (Table 2, Figure 3A). The problematic regions coincidentally aligned with relevant sgRNA regions (stem loop 2, repeat, and guide regions), potentially because of the RNA sequence (secondary structures, base composition, repeats) impacting the synthesis. High mismatch rates may reflect contamination with deprotected phosphoramidites during synthesis while high deletions may arise from incomplete coupling, capping, or oxidation reactions (Figure 1B). Correction of these regions was critical for achieving higher DNA cleavage activity using the sgRNA.

A particularly notable trend that occurred within all samples was high deletion rates within long homopolymers and high insertion rate for the site prior with their levels remaining relatively the same across all three samples. For instance, both the first U within the 3’-UUUU-5’ in the repeat/guide region and the first A within the 3’-AAAAA-5’ in stem loop 2 are long homopolymers that show a particularly high deletion rate. Furthermore, the A before 3’-UUUU-5’ as well as the G before 3’-AAAAA-5’ showed high insertions that were predominantly the same base type as the homopolymer (Figure 3A). The IVT 180mer similarly contained long homopolymers and similarly showed high deletion and insertion rates around the long homopolymer sites suggesting that the trend witnessed in the solid-phase synthesized samples could be due to an artifact from ONT DRS (Figure S1). During sequencing, homopolymers maintain the same current level, making them more difficult to distinguish compared to heteropolymers during basecalling, leading to higher deletion rates appearing during alignment to the reference [23].

The single-base analysis did not reveal any substantial error rates associated with base identity (Figure 4B). During basecalling, the most likely miscalls occur between guanine and adenine, both purines that produce similar current signals, and between cytosine and uracil, both pyrimidines. In contrast, miscalls between purines and pyrimidines are much less common [23]. Accordingly, the modestly elevated mismatches for U–C, C–U, and G–A observed in the confusion matrices (Figure 4C) are consistent with these expected, but infrequent, basecalling events. Unexpectedly, the first synthesis of GFP sgRNA showed elevated A-U mismatch levels that were higher than A-G. After optimization, A-U mismatch was reduced, suggesting there originally could have been synthesis issues related to the phosphoramidite reagents.

Our pipeline successfully leveraged ONT DRS to sequence truncated byproducts (Figure 2, S2-S4). The byproduct alignments to the reference were similar for all samples (first synthesis unmodified RNA and optimized synthesis unmodified and modified sgRNA). Reads within the <40 nt subset showed a bimodal distribution that could also be caused by an increase in coupling issues when nearing ∼50 nt lengths. This is similarly witnessed within the 40 – 60 nt subset where there are high occurrences of 50 nt length reads. The CPG pore size could be a potential limiter to RNA length during synthesis [36]. A 500 Å pore size can typically yield a maximum RNA length between 30 – 50 nt. Increasing pore size can promote longer RNA. However, it would decrease CPG surface area, leading to a reduction in synthesis yield. This drawback is one of many reasons why scaling-up RNA synthesis is so difficult and why many RNA synthesis companies either limits RNA quantity or length (i.e., high yield for short RNA or low yield for long RNA) [37–39].

Surprisingly, the proportion of full-length reads decreased after optimization: 9.10% for the initial synthesis, 7.04% for the unmodified optimized sample, and 3.89% for the modified optimized sample. Despite the lower proportion, previous analyses (Figure 3B) confirmed that the full-length reads generated after optimization were significantly more accurate, suggesting that the refined synthesis improved fidelity at the expense of yield.

Our results show how promising ONT DRS is for identifying RNA modifications. Ψ and m^6^A detection showed high confidence within their expected sites and were quantifiable within IGV. Surprisingly, the Ψ site showed significant amounts of U-C mismatch, suggesting that ONT’s SQK-RNA004 chemistry still results in systematic errors for Ψ sites.

Nm modified sgRNA results showed low to undetectable Nm sites. Nm is a common RNA modification used for sgRNA to improve the RNA stability. It is normally incorporated for the entire sgRNA sequence except for the guide region. Recently, the newest Dorado model (rna004_130bps_sup@v5.2.0) was released that added Nm to their list of detectable modifications. This allowed us to assess our 101 nt sgRNA with Nm modifications incorporated into the scaffold region, bringing about 58% of methylated bases overall (Table S1). The sequence visualization in IGV showed that the heavily Nm modified region was not basecalled correctly, leading to low coverage, low read count, and inability to detect the Nm sites (Figure S5 top). To enable accurate Nm detection in such contexts, further improvements will likely be required. These include: (i) training basecalling models on sgRNAs or other transcripts with dense Nm incorporation to improve recognition of modification-induced signal patterns, and (ii) developing analytical pipelines that integrate raw signal features with alignment-aware error models to distinguish modification-driven signatures from sequencing artifacts. Together, these advances could allow reliable basecalling and modification calling even in synthetic constructs with high densities of Nm.

We inspected the raw current signals of two representative regions with modification densities of 63% (Figure S5 bottom left) and 79% (Figure S5 bottom right). Surprisingly, both regions produced signals largely indistinguishable from the unmodified control, although certain stretches displayed clear current shifts. These observations suggest that very high Nm density may obscure the signal differences introduced by individual modifications, effectively masking their detection. Nonetheless, the fact that ONT DRS was able to register distinct signals in portions of these heavily modified sequences demonstrates a nascent capacity to resolve Nm in challenging contexts. With continued refinement of basecalling algorithms and training on highly modified RNAs, ONT DRS is poised to evolve into a powerful tool for direct detection of Nm even in densely substituted constructs.

## Conclusion

Incorporating ONT DRS into the RNA chemical synthesis pipeline led to improved product quality and provided insights unattainable with standard QC methods. This approach not only identified problematic regions arising during synthesis but also distinguished the types of errors and their nucleotide dependencies, enabling a deeper understanding of synthesis fidelity. ONT DRS showed distinct capabilities in identifying modification sites and with further refinement, the tool holds promise for resolving even densely modified RNA. In addition, extending this strategy to systematically analyze truncations will inform the causes of byproduct formation and support strategies to reduce them. Overall, our approach showed improved RNA sequence accuracy and biological efficacy that would not be achievable with only using the standard quality control methods.

## Data Availability

The sequencing data generated in this study have been deposited in the Sequence Read Archive (SRA) under accession code PRJNA1305499.

## Software Availability

The software tools and code to reproduce the analyses and plots are available on https://github.com/wanunulab/PRECISE-QC_analysis/tree/main.

## Supplementary Data

**Table S1.**
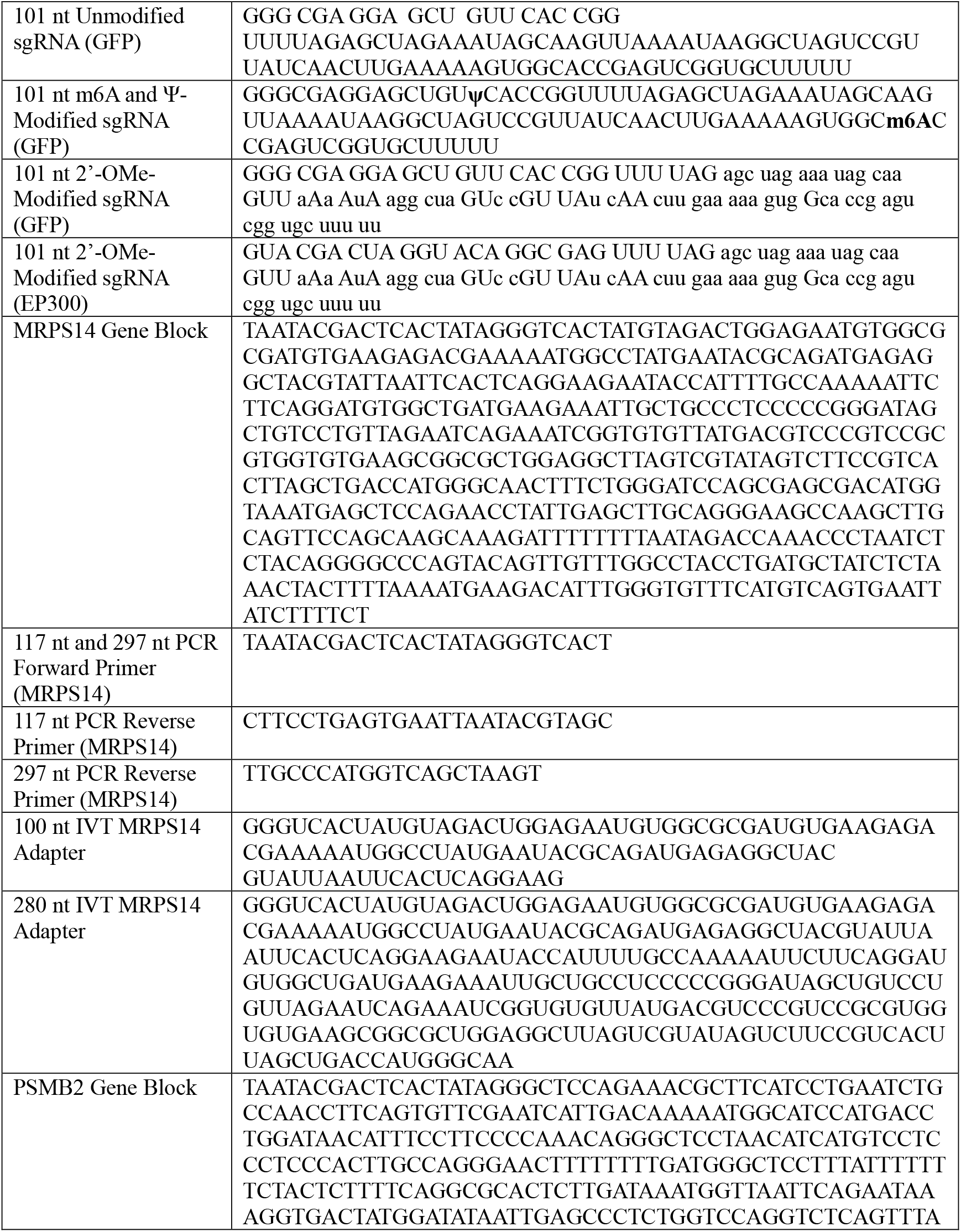
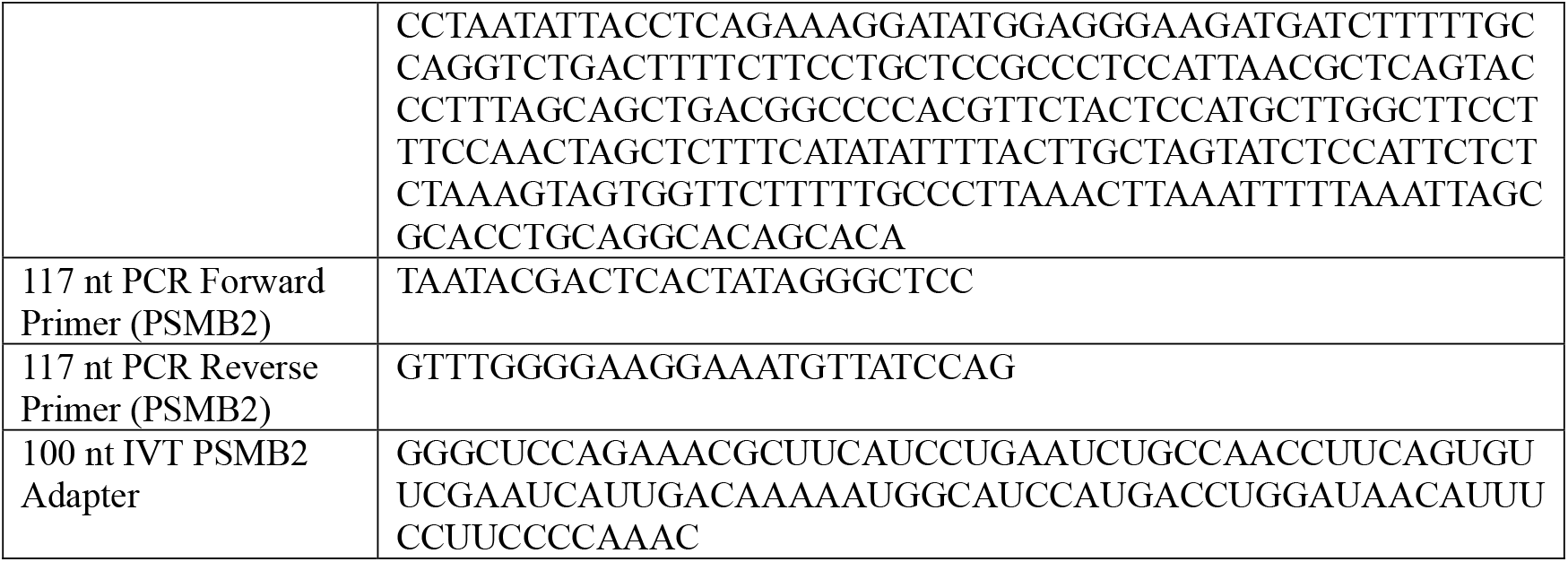
Sequences for all DNA and RNA used within the paper. All sequences are written 5’ to 3’. Capitalized letters are canonical bases while lowercase letters are 2’O methylated bases. Only 180 nt of the 280 nt IVT MRPS14 was used during the control analysis due to the 100 nt on the 5’ end matching the 100 nt adapter sequence complicating the results.

**Figure S1.**
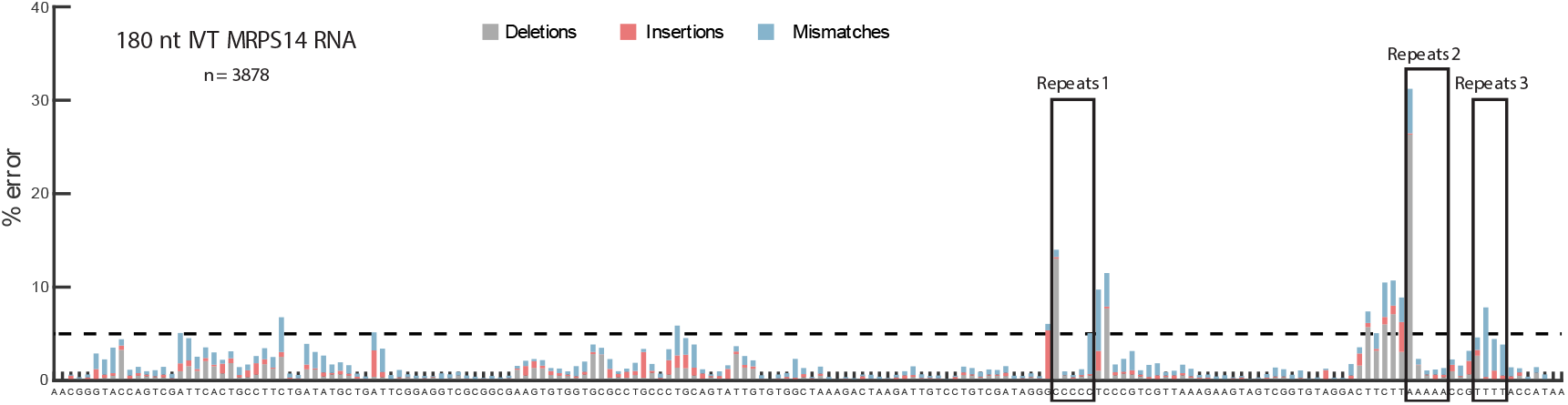
Error profile of an IVT synthesized 180 nt RNA. The dotted line sets the 5% error threshold to maintain above 90% synthesis accuracy. Boxed regions contain long homopolymers (>3 nt) showing high deletion rates for the first base of the sequence and high insertion rates for the base prior to the start of the homopolymer sequence.

**Figure S2.**
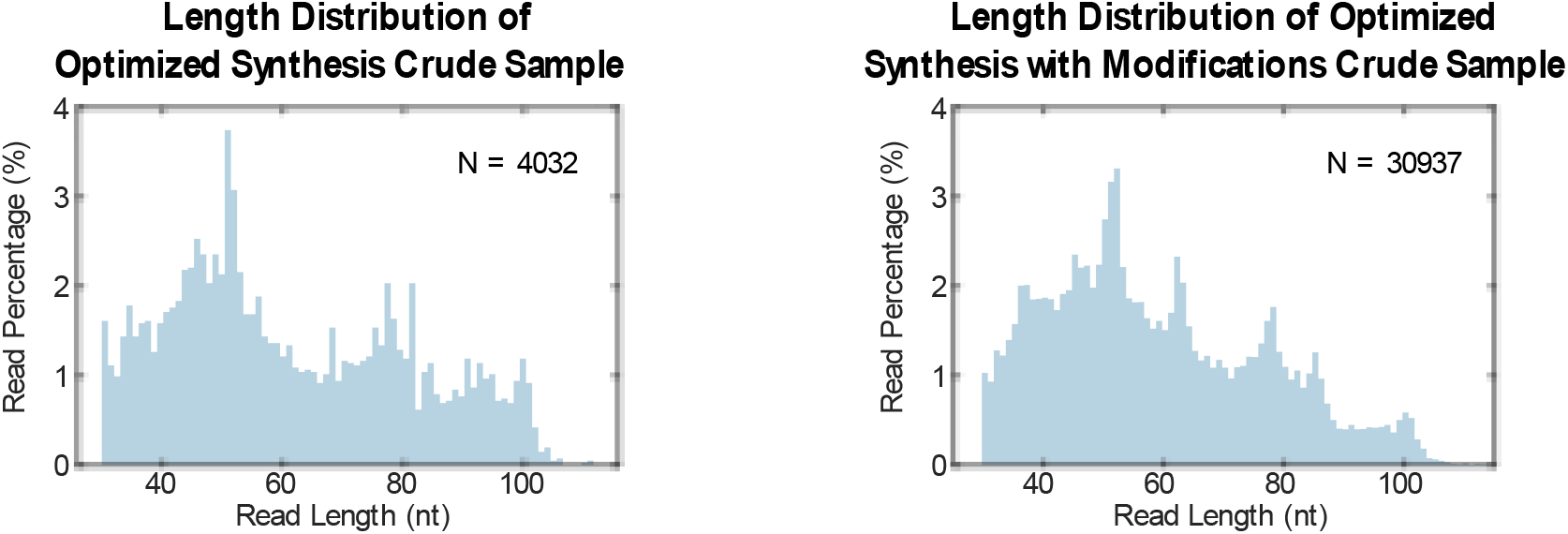
Length distribution of optimized synthesis crude samples.

**Figure S3.**
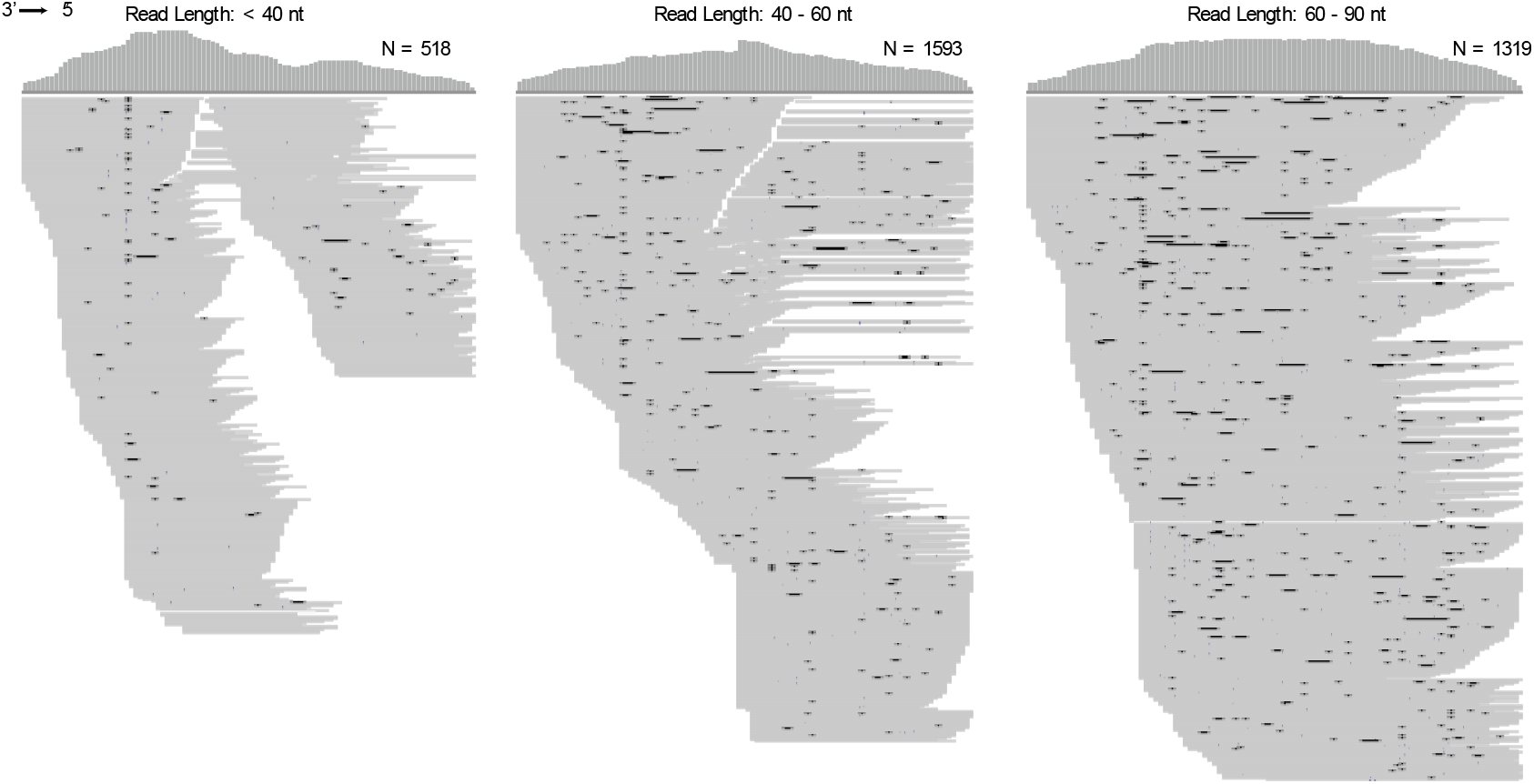
IGV and coverage plots for the unmodified GFP sgRNA optimized synthesis.

**Figure S4.**
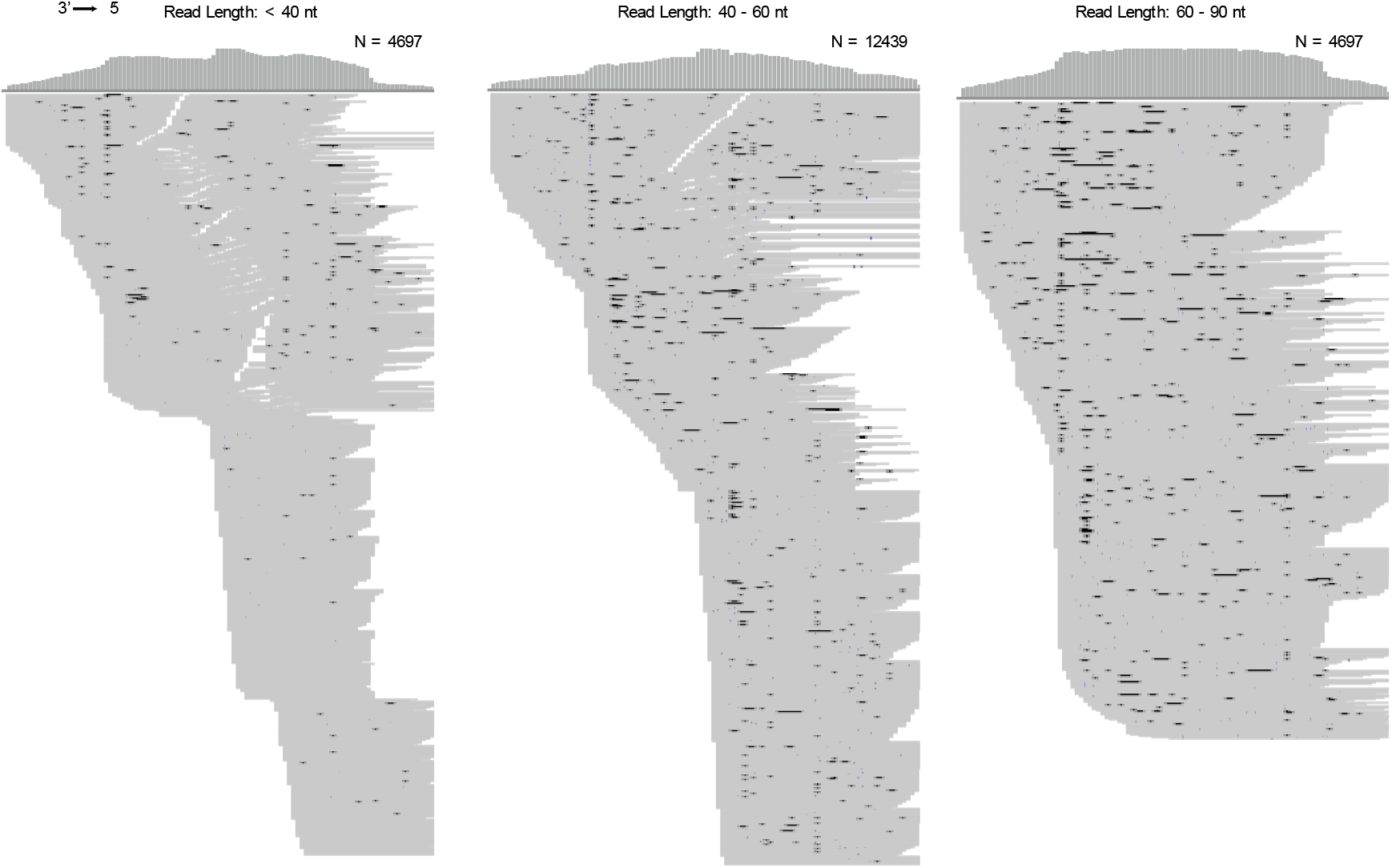
IGV and coverage plots for the modified GFP sgRNA optimized synthesis. Downsampling was used for easier visualization.

**Figure S5.**
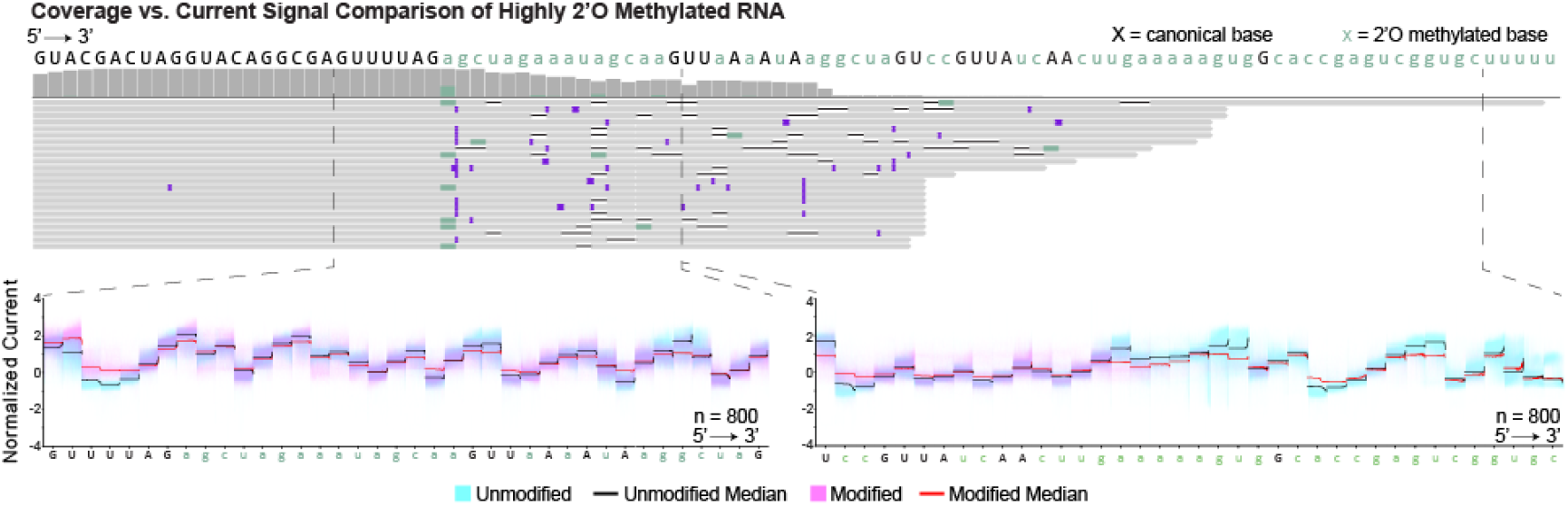
Comparison of IGV plot and coverage of highly Nm modified RNA to raw current signals at select regions.

## Author Contributions

BP and MR synthesized the sgRNA samples used in the study. YYY conducted the sequencing experiments. YYY and DB contributed to data acquisition and bioinformatic analyses. YYY, DB, SHR, MR, and MW conceived of and wrote the manuscript.

## Acknowledgements

We thank Miten Jain and Stuart Akeson for providing access to their GridION and Nanopore Direct RNA sequencing kit. We would like to further thank them for their helpful discussions regarding nanopore sequencing data analysis. We thank Parker Hitt for their artistic advice and assistance with figure graphics.

## Funding

The authors acknowledge generous support from the National Institutes of Health (R01HG013304 and R01HG012856).

## Conflict of Interest

The authors declare no competing interests.

## Notes

https://www.ncbi.nlm.nih.gov/sra/PRJNA1305499

### Competing Interest Statement

The authors have declared no competing interest.

