## Supplementary material for "Pore-Based RNA Evaluation for Control of Integrity, Sequence, and Errors – Quality Control (PRECISE-QC)": SI

#### Table 1

| | |
|---|---|
| 101 nt Unmodified sgRNA (GFP) | GGGCGAGGAGCUGUUCACCGGUUUUAGAGCUAGAAAUAGCAAGUUAAAAUAA GGCUAGUCCGU UAUCAACUUGAAAAAGUGGCACCGAGUCGGUGCUUUUU |
| 101 nt m6A and Ψ Modified sgRNA (GFP) | GGGCGAGGAGCUGU**ψ**CACCGGUUUUAGAGCUAGAAAUAGCAAGUUAAAAUAA GGCUAGUCCGUUAUCAACUUGAAAAAGUGGC**m6A**CCGAGUCGGUGCUUUUU |
| 101 nt 2'-OMe Modified sgRNA (GFP) | GGG CGA GGA GCU GUU CAC CGG UUU UAG agc uag aaa uag caa GUU aAa AuA agg cua GUc cGU UAu cAA cuu gaa aaa gug Gca ccg agu cgg ugc uuu uu |
| 101 nt 2'-OMe Modified sgRNA (EP300) | GUA CGA CUA GGU ACA GGC GAG UUU UAG agc uag aaa uag caa GUU aAa AuA agg cua GUc cGU UAu cAA cuu gaa aaa gug Gca ccg agu cgg ugc uuu uu |
| MRPS14 Gene Block | TAATACGACTCACTATAGGGTCACTATGTAGACTGGAGAATGTGGCGCGATGTGAAG AGACGAAAAATGGCCTATGAATACGCAGATGAGAGGCTACGTATTAATTCACTCAGG AAGAATACCATTTTGCCAAAAATTCTTCAGGATGTGGCTGATGAAGAAATTGCTGCC CTCCCCCGGGATAGCTGTCCTGTTAGAATCAGAAATCGGTGTGTTATGACGTCCCGTC CGCGTGGTGTGAAGCGGCGCTGGAGGCTTAGTCGTATAGTCTTCCGTCACTTAGCTG ACCATGGGCAACTTTCTGGGATCCAGCGAGCGACATGGTAAatgagctccagaacctattg agcttgcagggaagccaagcttgcagttccagcaagcaaagattttttttaatagaccaaaccctaatctctac aggggcccagtacagttgtttggcctacctgatgctatctctaaactacttttaaaatgaagacatttgggtgttt catgtcagtgaattatcttttct |
| 117 nt PCR Forward Primer (MRPS14) | TAATACGACTCACTATAGGGTCACT |
| 117 nt PCR Reverse Primer (MRPS14) | CTTCCTGAGTGAATTAATACGTAGC |
| 100 nt IVT MRPS14 Adapter | GGGUCACUAUGUAGACUGGAGAAUGUGGCGCGAUGUGAAGAGACGAAAAAUG GCCUAUGAAUACGCAGAUGAGAGGCUAC GUAUUAAUUCACUCAGGAAG |

*Supplementary Data*

| | |
|---|---|
| PSMB2 Gene Block | TAATACGACTCACTATAGGGCTCCAGAAACGCTTCATCCTGAATCTGCCAACCTTCAG TGTTCGAATCATTGACAAAAATGGCATCCATGACCTGGATAACATTTCCTTCCCCAAA CAGGGCTCCTAACATCATGTCCTCCCTCCCACTTGCCAGGGAACTTTTTTTTGATGGG CTCCTTTATTTTTTTCTACTCTTTTCAGGCGCACTCTTGATAAATGGTTAATTCAGAATA AAGGTGACTATGGATATAATTGAGCCCTCTGGTCCAGGTCTCAGTTTACCTAATATTAC CTCAGAAAGGATATGGAGGGAAGATGATCTTTTTGCCAGGTCTGACTTTTCTTCCTG CTCCGCCCTCCATTAACGCTCAGTACCCTTTAGCAGCTGACGGCCCCACGTTCTACTC CATGCTTGGCTTCCTTTCCAACTAGCTCTTTCATATATTTTACTTGCTAGTATCTCCATT CTCTCTAAAGTAGTGGTTCTTTTTGCCCTTAAACTTAAATTTTTAAATTAGCGCACCTG CAGGCACAGCACA |
| 117 nt PCR Forward Primer (PSMB2) | TAATACGACTCACTATAGGGCTCC |
| 117 nt PCR Reverse Primer (PSMB2) | GTTTGGGGAAGGAAATGTTATCCAG |
| 100 nt IVT PSMB2 Adapter | GGGCUCCAGAAACGCUUCAUCCUGAAUCUGCCAACCUUCAGUGUUCGAAUCAU UGACAAAAAUGGCAUCCAUGACCUGGAUAACAUUUCCUUCCCCAAAC |

Capitalized letters are unmodified bases while lowercase letters are 2’-OMe

#### Figure S1

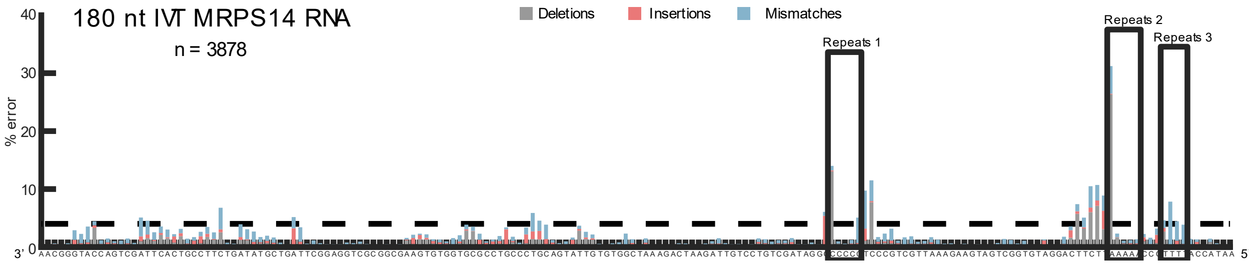


Error profile of an IVT synthesized 180 nt RNA. Boxed regions contain long homopolymers (> 3 nt) showing high deletion rates for the first base of the sequence and high mismatch rates for the base prior to the start of the homopolymer sequence.

### Figure S2

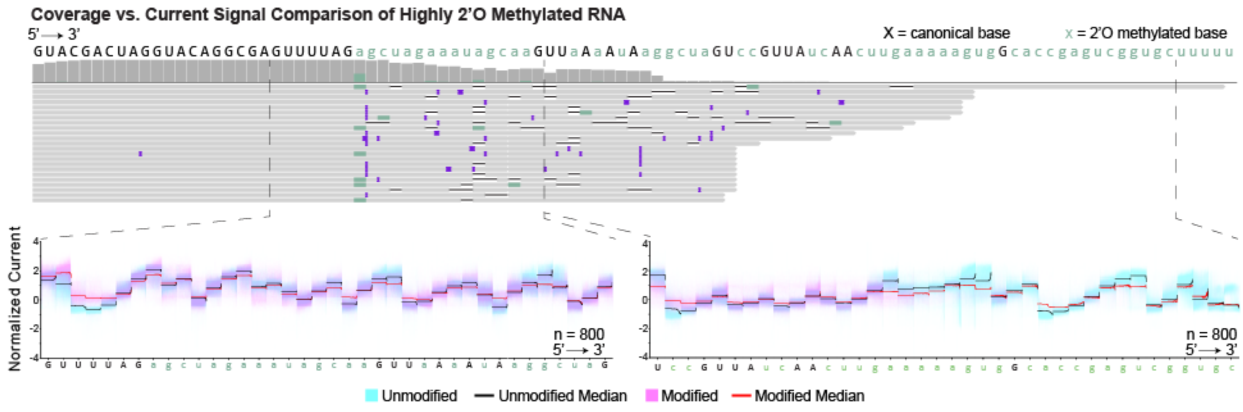
Coverage vs. Current Signal Comparison of Highly 2'O Methylated RNA
5' → 3'
X = canonical base
x = 2'O methylated base
Normalized Current
n = 800
5' → 3'
n = 800
5' → 3'
Unmodified
Unmodified Median
Modified
Modified Median
